# Low-dose infusion of a superagonistic anti-CD28 monoclonal antibody ameliorates inflammation after acute myocardial infarction in pigs

**DOI:** 10.1101/2025.10.28.685145

**Authors:** Anja Stadtmüller, Rebekka Grampp, Florian Schnitter, Finja Becker, Könül Nuriyeva, Daniela Langenhorst, Leona Metka, Kira Günther, Nadine Gladow, Ulrich Hofmann, Anna Frey, Niklas Beyersdorf

## Abstract

CD4^+^ Foxp3^+^ regulatory T cells (Treg) efficiently foster wound healing after myocardial infarction (MI). Moreover, therapeutic shifting of the balance between CD4^+^ Foxp3^−^ conventional (Tconv) and Treg towards Treg enhances survival in mice after MI. Owing to the species-specific differences in cardiac wound healing and remodelling after MI, it remains unclear whether these findings can be translated into novel immunotherapies for patients with MI. Therefore, we tested the feasibility of therapeutic Treg activation after MI in pigs whose cardiac wound healing and the composition of the immune system are very close to humans. To activate Treg, we treated pigs three days after MI induction with a superagonistic anti-CD28 monoclonal antibody (mAb, CD28-SA; clone 4D12). To avoid induction of a cytokine release syndrome (CRS) we infused the CD28-SA at a low dose of 10 µg/kg body weight. Low-dose CD28-SA treatment led to a local increase in porcine Treg among heart-infiltrating CD4^+^ T cells in comparison with the control mAb therapy (clone MOPC-21). Immunofluorescence histology showed that CD28-SA application particularly strengthened Treg accumulation in the infarct border zone. This was paralleled by a lower proportion of TNFα-producing cells among cardiac monocytes and macrophages reflecting a shift away from inflammation and towards wound healing. Taken together, our findings in the pig MI model are the first to show that low-dose CD28-SA infusion is capable of ameliorating tissue inflammation in a species prone to CRS development after high-dose CD28-SA infusion. Moreover, the data highlight that therapeutically shifting the Treg/Tconv balance in MI patients might be feasible in order to enhance cardiac wound healing and prevent adverse remodelling.

## Introduction

Myocardial infarction (MI) has a high prevalence and, when survived, often leads to the development of chronic heart failure which by itself comes with a high mortality (1). Whereas the therapeutic gold standard for MI aims for rapid restitution of coronary blood flow (1), unfortunately only 54% of patients undergo timely reperfusion (2). The remaining patients often suffer extensive transmural infarction, resulting in tissue necrosis and subsequent adverse myocardial remodelling (3). However, there are currently no therapies available to therapeutically modulate cardiac wound healing after MI and to prevent adverse remodelling despite the considerable efforts that have been made to change this (4). Targeting cardiac inflammation post MI has the potential for a breakthrough in this field as inflammation is one of the driving forces behind early healing as well as mid- and long-term remodelling processes (5). In mouse and rat models of myocardial infarction, we and others were able to establish that therapeutically shifting the balance of CD4^+^ Foxp3^−^ conventional (Tconv) and CD4^+^ Foxp3^+^ regulatory T cells (Treg) towards Treg enhances wound healing, cardiac remodelling and long-term survival after MI (6, 7). A shift in the Treg/Tconv balance towards Treg could either be achieved with a superagonistic anti-CD28 monoclonal antibody (CD28-SA) (7) or by injection of low-dose interleukin 2 (IL-2) (8, 9). Alternatively, blocking T cell costimulation with an anti-CD28 mAb blocking ligand binding (10) or with CTLA-4-Ig (Abatacept) (11) preferentially inhibited Tconv over Treg also leading to a shift in the Treg/Tconv balance towards Treg and a better outcome after MI.

Compared to laboratory mice and rats, the human immune system faces many more challenges leading to ample formation of memory cells in the T and B cell compartments and a state of ‘training’ in cells of innate immunity (12). These differences pose significant challenges for the transfer of immunotherapeutic approaches from rodents to humans. In case of MI, the sheer size of the human heart is another important difference to mice and rats, leading to markedly slower wound healing (13). Therefore, we assessed whether Treg activation after MI was feasible in conventionally housed pigs, i.e. in a human-like animal model. For Treg activation we used a CD28-SA which we had shown to have the same dose-response relationship *in vivo* as the human CD28-SA TAB08 (formerly TGN1412) (14). As for IL-2 infusion, CD28-SA treatment also comes with the risk that high dosages are capable of inducing life-threatening toxicities, which in the case of CD28-SA are due to the induction of a cytokine release syndrome (CRS).

In this study, we thus infused the CD28-SA into pigs after MI at a low dose of 10 µg/kg bw (body weight). In-depth cardiological and immunological assessment of these animals until day seven post MI revealed, on the one hand, no negative effects of CD28-SA infusion on cardiac function, while, on the other hand, CD28-SA increased the proportion of Treg among CD4^+^ T cells in the infarcted heart and reduced the proportion of TNFα-producing cells among cardiac monocytes and macrophages.

## Material and Methods

### Animals and ethics statement

Pigs (German Landrace) were bred on Gerd Heinrichs’ farm (Heinsberg, Germany) and maintained in the animal facility of the Comprehensive Heart Failure Center Würzburg at the University Hospital of Würzburg. All animal experiments were conducted in accordance with the Animal Protection Law (Directive of the European Parliament and of the Council of 22 September 2010 (2010/63/EU)) and had been reviewed and approved by the review board of the District Government of Lower Franconia and the University of Würzburg (approval reference number 55.2.2.-2532-2-1402). The experiments were carried out in accordance with the ethical principles of the Declaration of Helsinki (1964, and later amendments) and the guidelines of the Federation of European Laboratory Animal Science Associations (FELASA). Surgical procedures were performed under general anaesthesia, and appropriate analgesia was provided to alleviate postoperative pain. Additionally, all efforts were made to minimise animal suffering and to reduce the number of animals used.

### Pig MI model

Pigs (aged 10 to 20 weeks with an average bw of 43 ± 10 kg) underwent MI induction under total intravenous anaesthesia with propofol (5 mg/kg bw/h i.v.) and fentanyl (12-24 µg/kg bw/h i.v.) according to the established protocol in our group (15). In brief, pigs were sedated with azaperone (8 mg/kg bw i.m.), including atropine (0.5 mg), anaesthesized with ketamine (15 mg/kg bw i.m.), intubated after a propofol bolus (2 mg/kg bw i.v.) and put on a mechanical ventilation for the duration of the procedure. After femoral artery puncture and subsequent coronary catheterisation, the pigs underwent temporary balloon occlusion of the left anterior descending artery (LAD) for 90 minutes (Supplemental Figure S1). Four pigs died during the LAD occlusion due to electrotherapy-resistant ventricular fibrillation despite continuous application of amiodarone (2 mg/kg bw/h i.v.) for arrhythmia prophylaxis. For additional pain prevention post-MI, the pigs received meloxicam (0.4 mg/kg bw i.m.) before completion of the anaesthesia at the end of the procedure and on the following day.

### Echocardiography

To analyse cardiac function, transthoracic echocardiography was performed on day 0 before MI (baseline), as well as on day 3 and on day 7 before termination of the experiment with an ultrasound system (Vivid E9; GE Healtcare, Chicago, IL, USA) under anaesthesia. Echocardiography was first performed from the right subxiphoid view (5-chamber), followed by left parasternal long- and short-axis views first in left and then in right lateral position. Parameters to assess cardiac function included stroke volume (LVOT diameter and curve), systolic and diastolic function (left ventricular ejection fraction using the Teichholz method, M-mode, E/A, E/É), and wall motion. Some recordings were acquired during short breath-holds under mechanical ventilation to improve image quality, which was not feasible in spontaneously breathing animals. The evaluation of parameters was performed in a post-processing manner by a blinded doctoral researcher. All parameters were measured three times for each animal and the arithmetic mean was used for comparative analysis between the groups.

### Antibody infusion

On day 3 after MI, pigs were again sedated and intubated as described above, but now subjected to inhalation anaesthesia with isoflurane (1-2 Vol. %) in order to maintain spontaneous breathing and facilitate later recovery. In advance, the animals had independently and randomly been assigned to either the treatment group receiving therapeutic CD28-SA (clone 4D12, InVivo BioTech Services GmbH, <1 EU/mg Endotoxin level), or to the control group receiving isotype control antibody at the same dosage of 10 µg/kg bw (MOPC-21, BioXCell, <1 EU/mg Endotoxin level). The agent was provided in a blinded manner in 10 ml 0.9% sterile saline solution and then further diluted in 500 ml 0.9% sterile saline. The total volume of 510 ml was administered i.v. within 30 minutes under continuous cardiopulmonary monitoring. All twelve pigs survived the antibody infusion and exhibited no signs or symptoms of an allergic reaction or even a CRS. The researchers who performed the MI, and the echo, applied the infusion and conducted the subsequent processing of the harvested organs were blinded to the antibody infused to obtain unbiased results. Unblinding for infused therapy occurred only after all reported individual measurements had been evaluated and the analysis of the whole data set had started.

### Serial blood sampling and preparation of PBMC

Blood samples were taken on day 0, i.e. before MI induction, as well as before the procedures on days 3 and 7. Serum samples were centrifuged, the supernatant was collected and preserved in aliquots at −80 °C. Furthermore, heparinised blood samples were taken for whole blood analysis and for peripheral blood mononuclear cell (PBMC) isolation via the Ficoll separation method (Histopaque-1077, Sigma-Aldrich). The isolated PBMC were resuspended in cell culture medium RPMI-1640 containing 10% fetal calf serum (FCS, Gibco), 2-Mercaptoethanol, sodium pyruvate, MEM solution, 5% Glutamin, Streptomycin, Penicillin and counted. 1×10^6^ cells (2×10^6^ on day 7) were further stained for flow cytometry and the remaining cells were preserved in cell cryomedium (RPMI-1640, FCS, Dimethylsulfoxid – DMSO) at −80 °C.

### Organ preparation on day seven post MI

On day 7 after MI anaesthesia was induced as described above. Blood samples were taken from the animals and the final echocardiography was completed. After bolus administration of heparin i.v. (300 IU/kg bw) pigs were euthanised with pentobarbital i.v. (dosage according to manufactureŕs instructions) for immediate organ extraction. Parts of the spleen, thymus and mediastinal lymph nodes were removed and submerged in cold RPMI-1640 medium. From there, the samples were cut into smaller pieces, weighed and grounded through a 70 μm cell strainer into cold Hankś Balanced Salt Solution without Ca^2+^ and Mg^2+^, with phenol red and with 0.1% bovine serum albumin (HBSS -/- /BSA). With the Ficoll separation method, splenocytes and thymocytes were isolated. After a final centrifugation, all the cell suspensions were resuspended in a cell culture medium and stored overnight at 4 °C. On the next day, the cells were counted and either further stained for flow cytometry or preserved in cell cryomedium.

After removal, the hearts were washed with ice-cold 0.9% saline. In addition, the coronaries on both sides were manually flushed to remove any blood residues. The organ was weighed and photographed. The left ventricle was then cut into evenly thick transverse slices (8-10 mm), which were weighed and photographed individually. Samples of 600 mg tissue from the different regions, i.e. infarct core (IC), border zone (BZ), and remote myocardium (RM), were taken from infarcted heart slices. At least two different sites were sampled per region - the respective regions had been selected purely macroscopically (Supplemental Figure S1). The specimens were next processed for immunofluorescence staining and flow cytometry.

To isolate leukocytes, the heart tissue was finely chopped and digested at 37 °C for 30 minutes with DNAse I (60 units/ml Roche Diagnostics GmbH; Mannheim, Germany) and collagenase type 2 (600 units/ml Worthington Biochemical, Lakewood, NJ, USA) in Hanks’ balanced salt solution (HBSS +/+, with Ca^2+^/Mg^2+^ without phenol red). The samples were then first passed through 70 µm cell strainers and then through 40 µm cell strainers in cold HBSS -/- /BSA. This was followed by 8 minutes of red blood cell lysis with an RBC-lysis buffer (10x, BioLegend, San Diego, CA, USA) and live-dead staining with Zombie Aqua 1:200 (fixable viability kit (DMSO); BioLegend, San Diego, CA, USA) at room temperature, darkened, for 15 minutes in DPBS−/− /EDTA (Dulbecco’s Balanced Salt Solution without Ca^2+^/Mg^2+^, containing 2 mM ethylene diamine tetra-acetic acid). After washing the samples with flow cytometry buffer (DPBS−/− /EDTA with 2 % FCS), the cells were blocked with pig serum 10 % (own production; in DPBS−/− /EDTA) for 15 minutes at 4 C. The next step was the anti-CD45-FITC antibody staining (1:50, Mouse anti Pig CD45 FITC; Bio-Rad; clone: K252.1E4) 30 minutes at 4 C. To ensure a sufficient yield of leukocytes from all three heart areas (IC, BZ, RM) magnetic activated cell sorting (MACS) was performed. For that, after washing the cells with flow cytometry buffer, they were incubated for 15 minutes at 4°C with anti-FITC MicroBeads (1:20, Miltenyi Biotec, Bergisch Gladbach, Germany). Another washing step followed and CD45^+^ cells were positively isolated using LS columns (Miltenyi Biotec, Bergisch Gladbach, Germany) according to the manufacturer’s instructions.

### Flow cytometry. Whole blood samples and PBMC

50μl of whole blood were transferred to TruCOUNT tubes containing TruCount Beads (BD Biosciences, Franklin Lakes, NJ, USA) and processed following the manufacturer’s instructions - including calculation of absolute cell numbers. Antibodies used for staining are listed in Supplemental Table 1. Whole blood samples were analysed on a FACS Calibur (BD FACSCalibur^TM^ Flow Cytometer, BD Biosciences). Isolated PBMC were stained with the antibody panel summarised in Supplemental Table 2. Supplemental Figure S2 shows an example of our gating strategy.

### Cardiac leukocytes

Antibodies used for staining cardiac leukocytes (lymphocytes and myeloid cells) are listed in Supplemental Tables 2 - 3. The fixation and permeabilisation kit (eBioscience Foxp3/Transcription Factor Staining Buffer Set - Thermo Fisher Scientifc, Carlsbad, CA, USA) was used to process the samples incubated with the lymphocyte antibody panel (see Supplemental Table 4). In addition to cells from the heart, cells from **mediastinal lymph nodes, spleen and PBMCs** were stained using the lymphocyte panel.

Data analysis was conducted using FlowJo v10.10.0 (FlowJo, Ashland, OR, USA) following measurements obtained with the Attune NxT Flow Cytometer (Thermo Fisher Scientific, Waltham, MA, USA) and Calibur.

### Detection of TNFα producers among cardiac myeloid cells

TNFα producers were detected among cells collected from the heart on day 7. A total of 2 × 10^5^ cells were resuspended and seeded into each well of a 96-U well plate as duplicates. To stimulate the production of cytokines, the cells were stimulated with a mixture of 500 ng/ml of ionomycin and 5 ng/ml of PMA. As a control, wells were left without any stimulation. After an incubation period of 2 hours at 37°C, 5% CO2, Brefeldin A (10 μg/ml) was added to the cultures and the cells incubated for an additional 2 hours. Following incubation, the contents of the sterile 96-U well plate were resuspended thoroughly and transferred to a non-sterile 96-V well plate. Centrifugation was performed at 4°C for 3 minutes at 1600 rpm to pellet the cells, and the cytokine staining was carried out according to Supplemental Table 3. Following staining, the cells were resuspended in 100 μl FACS buffer and analysed on a BD™ LSR II Flow Cytometer. Supplemental Figure S3 shows an example of our gating strategy.

### Immunofluorescence histology of cardiac tissue

For immunofluorescence staining samples of cardiac tissue from different regions (IC, BZ, RM) were embedded in cryomedium (Tissue-Tek; Sakura Finetek Europe, Alphen aan den Rijn, The Netherlands) on dry ice and preserved at −80 °C. A few days before staining, the frozen tissue was cut into 7 µm thick slices with a cryostat microtome (CM1850; Leica Biosystems, Wetzlar, Germany), applied to microscope slides and retained at −20 C. The cryosections were thawed for 30 minutes at room temperature and fixed in 4% PFA (paraformaldehyde). Then, the slides were washed in tris-buffered saline (TBS) 3 times for 5 minutes and permeabilized with 0.02% Triton (Triton X-100) for 10 minutes. Washing steps were repeated, followed by the application of the blocking medium (60 minutes – IHC/ICC Blocking Buffer High Protein, Invitrogen) in a humid chamber at room temperature. Afterwards, anti-CD4 PE (Mouse anti Pig CD4 Alpha RPE; Bio-Rad; clone MIL17) was applied (1:50 diluted in blocking medium) and the samples were stored at 4°C overnight (continuously in the humid camber). On the following day, the washing routine was changed to initiate the buffer switch [2 x 5 minutes in TBS and 1 x 5 minutes in phosphate-buffered saline (PBS)]. Next, the permeabilization with 0.5% dodecyltrimethylammonium-chloride (DTAC) for 20 minutes took place and was followed by staining with anti-Foxp3 APC (monoclonal antibody; eBioscience™; Invitrogen; clone FJK-16s) 1:200 diluted in blocking medium at 4 °C overnight in the humid chamber. Lastly, the slides were washed with PBS (3 x 5 minutes) and the cell nuclei were stained with DAPI (4′,6-diamidino-2-phenylindole – 1:5000) for 10 minutes at room temperature in the humid chamber. The washing steps were repeated and the samples were mounted with ImmuMount (Epredia Netherlands B.V., DA Breda, Netherlands). For analysis, the DMi8 fluorescence microscope (Leica Microsystems, Wetzlar, Germany) was used. The sections were examined manually using ImageJ. At least 100 CD4+ cells should be counted, but no more than 6 visual fields (715µm x 530µm) were considered. For the BZ definition, both the IC and the RM had to be visible in the selected field. CD4^+^ cells and double-positive CD4^+^ Foxp3^+^ cells were assessed separately. The cell counts were then put into relation to the number of counted visual fields.

### Statistical data analysis

Statistical analysis and data visualisation were performed using GraphPad Prism v10.6.0 (GraphPad Software, San Diego, CA, USA). Values for each animal are shown individually with horizontal bars indicating means per group. The number of samples available for each group and the statistical tests applied are specified in the figure legends. A p value < 0.05 was considered statistically significant.

## Results

### CD28-SA infusion did not affect the deterioration of cardiac function within the first week after MI

Both in the pig MI model (16) as well as in humans (1) with similarly large infarcts, cardiac function continues to deteriorate in the days following MI. To determine whether low-dose CD28-SA infusion (10 µg/kg bw) has a short-term impact on cardiac function, we conducted serial echocardiography on pigs before and after MI induction. For these analyses, we included data from a total of twelve pigs (six treated with the CD28-SA clone 4D12 and six treated with the control mAb MOPC) that survived the whole observation period. In echocardiography, reductions in the left ventricular ejection fraction (LV-EF, Figure 1D) and the fractional shortening (LV-FS, Figure 1E) clearly indicated the expected deterioration in cardiac function during the first week after MI. In addition, disturbed motion of both the septal and the anterior walls of the left ventricle confirmed successful induction of MI (Supplemental Videos). However, there were no statistically significant differences between CD28-SA-treated pigs and pigs that had received the control mAb MOPC in any of the parameters analysed (Figure 1). The echocardiographic data, thus, showed that low-dose CD28-SA infusion had no negative side-effects on cardiac function in any of the infarcted pigs. The short observation period and the early timepoints after MI analysed might have precluded observing positive effects of CD28-SA infusion on cardiac remodelling in a long-term perspective as previously observed in mice (7).

**Figure 1.**
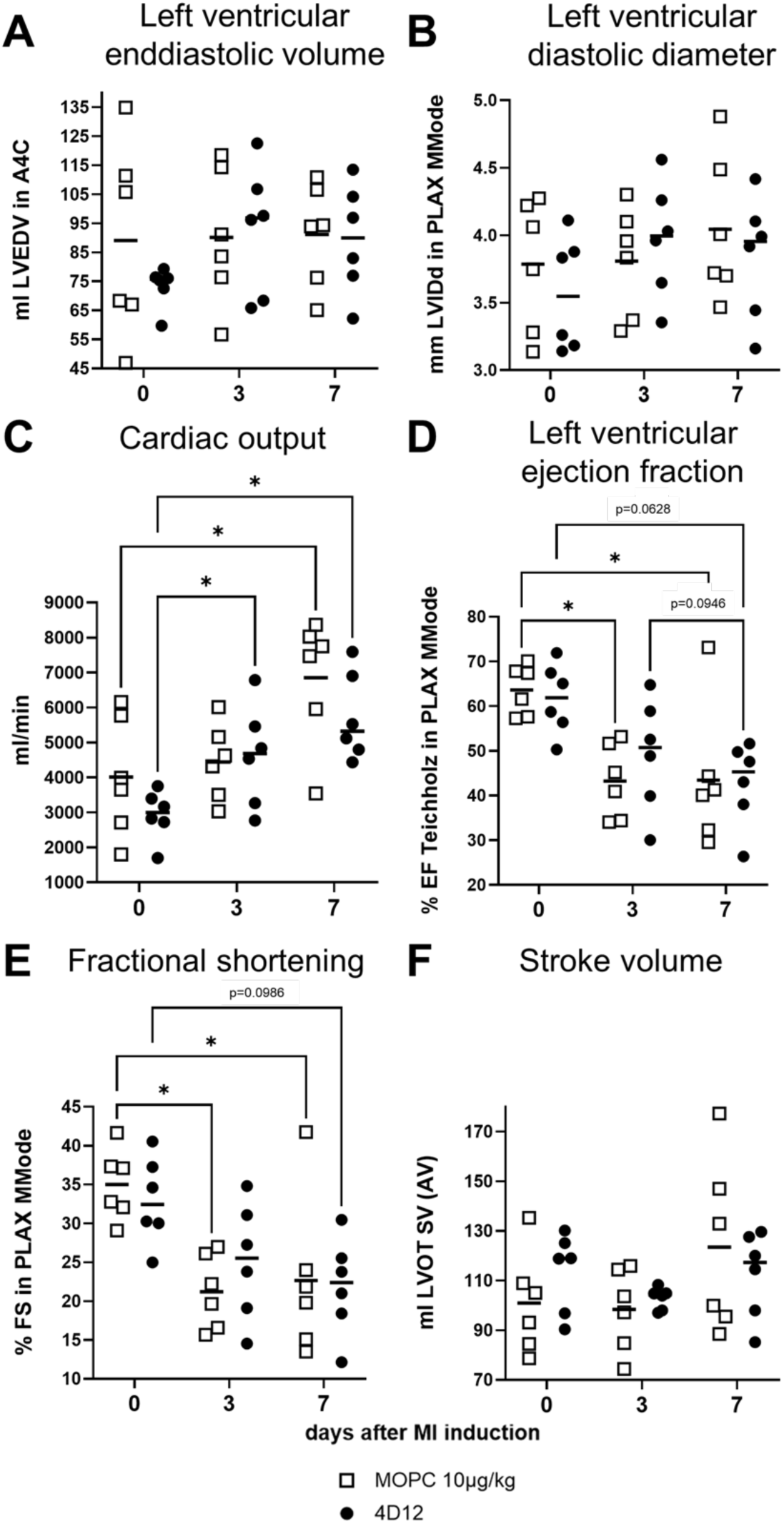
Similar cardiac function as determined by echocardiography in pigs on day 7 after MI irrespective of antibody treatment. **A**) Left ventricular enddiastolic volume (LVEDV) in the apical 4-chamber view (A4C) is stable over time after MI without differences between both groups. **B**) Left ventricular internal diastolic diameter (LVIDd) in parasternal long-axis view (PLAX) is stable over time after MI without differences between both groups. **C**) Cardiac output is increasing over time without differences between both groups due to the growth of the animals. **D**) Left ventricular ejection fraction (EF) by the Teichholz method in PLAX is decreasing over time without differences between both groups due to the MI induction. **E**) Fractional shortening (FS) in PLAX is decreasing over time without differences between both groups due to MI induction. **F**) Stroke volume (SV) by the single-plane Simpson’s method of discs in A4C is stable over time after MI without differences between both groups. Stats: Repeated measures two-way ANOVA followed by Tukey’s post-hoc test. 0.05 ≤ p < 0.1 are shown as numbers. * p < 0.05, ** p < 0.01.

### Detection of biologically active CD28-SA in peripheral blood and secondary lymphoid organs

In our earlier study we compared the effect of infusion of low-dose CD28-SA versus control mAb MOPC into healthy pigs (14). However, with three pigs per group the data were not conclusive as to how CD28-SA infusion affected the T cell compartment in serial analyses of peripheral blood.

In the infarcted animals, the CD28-SA (clone 4D12) infused on day three after MI induction was readily detectable on CD4^+^ T cells from peripheral blood (Figure 2A), spleen and mediastinal lymph nodes (Figure 3A) on day seven after MI induction. As a consequence of 4D12 infusion, CD28 expression was increased on CD4^+^ T cells (Figure 2B, 3B). Neither MI induction nor antibody infusion had an impact on the absolute numbers of total T cells or of CD4^+^ and CD8^+^ T cells in peripheral blood (Figure 2C - E). While there were no differences between 4D12- and MOPC-treated animals regarding the proportion of CD4^+^ T cells among leukocytes of spleen (Figure 3C), mediastinal lymph nodes (Figure 3C) and blood (Figure 2F), the proportion of Treg among CD4^+^ T cells was increased in peripheral blood on day seven both in 4D12- and MOPC-treated animals. (Figure 2G). As for peripheral blood, Treg frequencies among CD4^+^ T cells of spleen and mediastinal lymph nodes were not increased after 4D12 compared to MOPC infusion (Figure 3D). The proportion of proliferating Ki-67^+^ cells among Treg of peripheral blood (Figure 2H), spleen and mediastinal lymph nodes (Figure 3E) were also similar in 4D12- and MOPC-treated pigs. However, for CD4^+^ CD25^−^ Foxp3^−^ Tconv of peripheral blood we observed an increased proportion of Ki-67^+^ cells after 4D12 compared to MOPC infusion (Figure 2I), which was not the case in spleen and mediastinal lymph nodes (Figure 3F). The CD28-SA 4D12 could thus be readily detected on CD4^+^ T cells four days after the infusion, inducing an increase in CD28 expression in all three compartments as well as a higher proportion of proliferating cells among blood Tconv cells.

**Figure 2.**
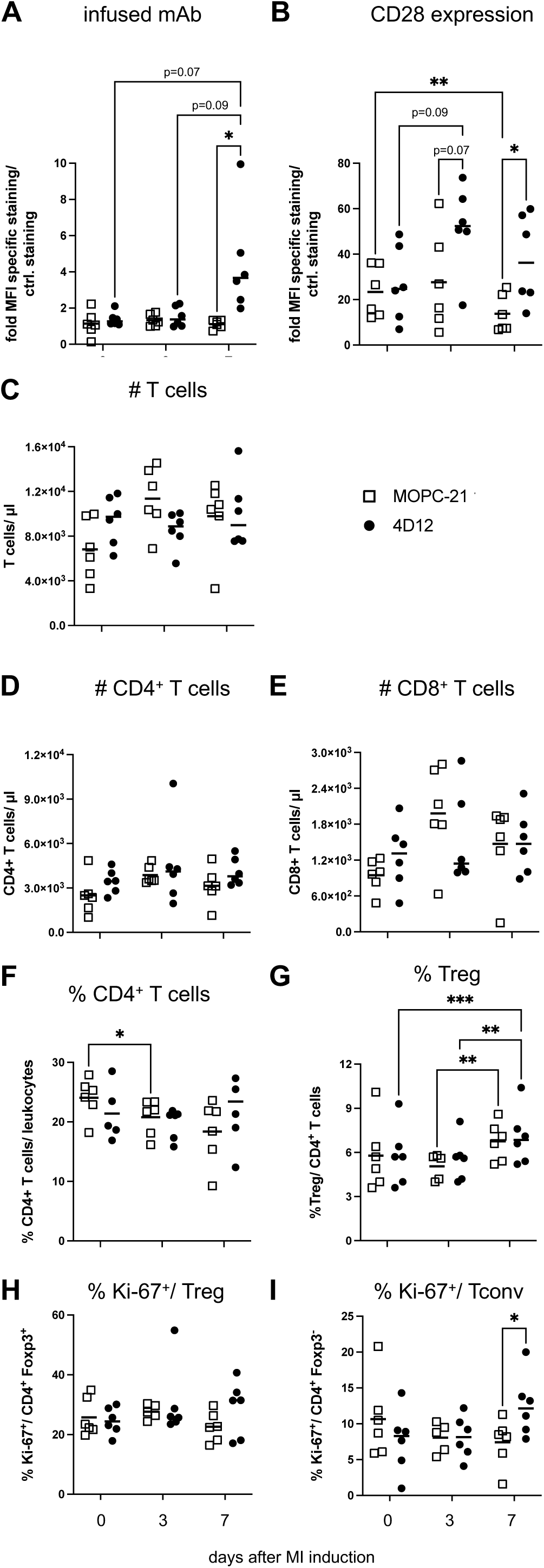
Tracking of infused CD28- SA *in vivo* and longitudinal analysis of T cell subsets in peripheral blood. **A)** Detection of mAb 4D12 on CD4^+^ T cells of peripheral blood. **B**) CD28 expression by peripheral-blood CD4^+^ T cells. **C**) Absolute number of total T cells per μl of blood. **F**) Proportion of CD4^+^ T cells among peripheral-blood leukocytes. **G**) Proportion of CD25^+^ Foxp3^+^ Treg among CD4^+^ T cells. **H**) Proportion of Ki-67^+^ cells among Treg and **I**) Tconv. Stats: Repeated measures two-way ANOVA followed by Tukey’s post-hoc test. 0.05 ≤ p < 0.1 are shown as numbers. * p < 0.05, ** p < 0.01, *** p < 0.001.

**Figure 3.**
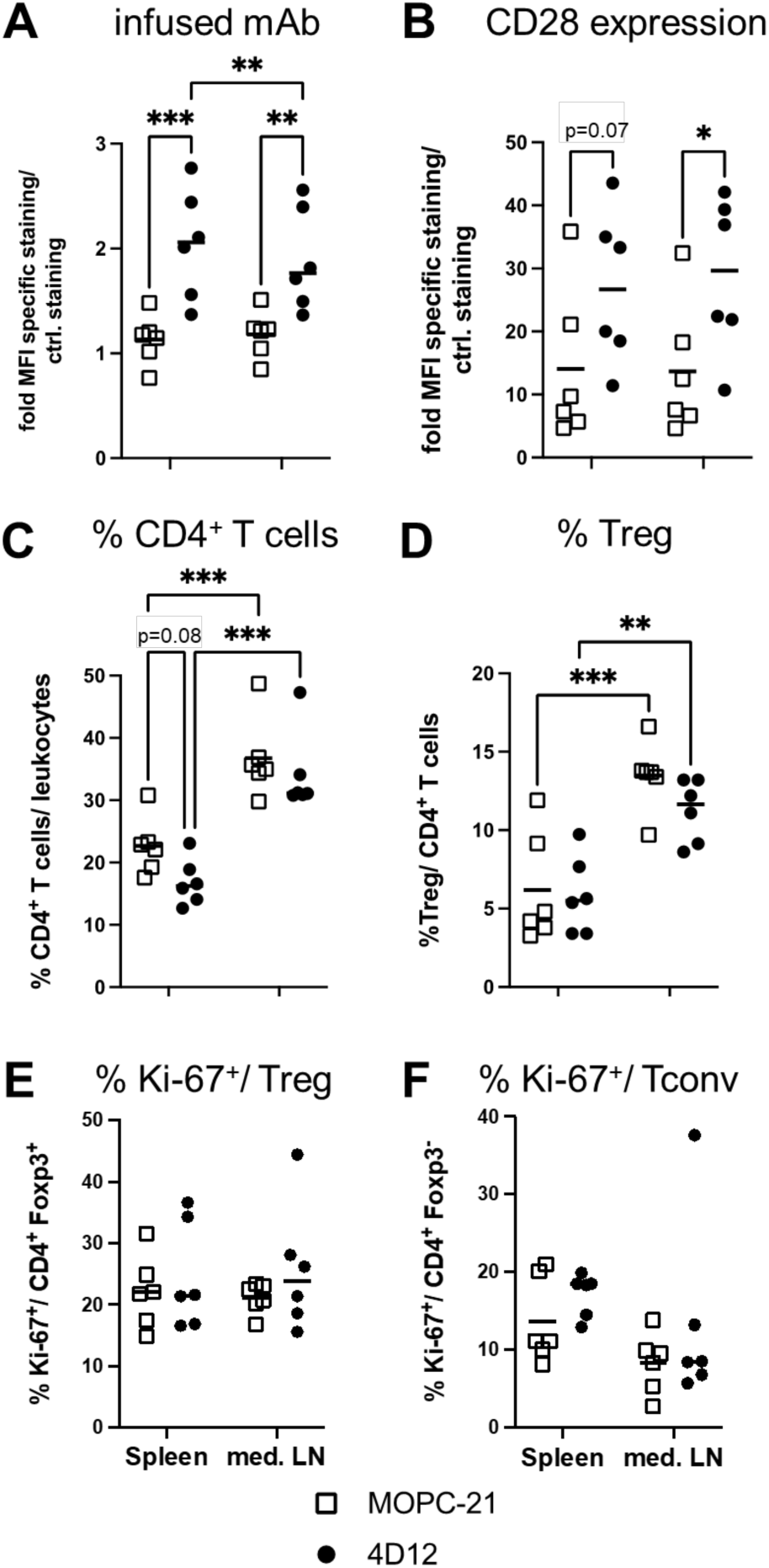
CD28-SA infusion increased CD28 expression on CD4^+^ T cells of the spleen and mediastinal lymph nodes on day 7 after MI. **A**) Detection of mAb 4D12 on CD4^+^ T cells. **B**) CD28 expression by CD4^+^ T cells. **C**) Proportion of CD4^+^ T cells among leukocytes. **D**) Proportion of CD25^+^ Foxp3^+^ Treg among CD4^+^ T cells. **E**) Proportion of Ki-67^+^ cells among Treg and **F**) Tconv. Stats: Repeated measures two-way ANOVA followed by Tukey’s post-hoc test. 0.05 ≤ p < 0.1 are shown as numbers. * p < 0.05, ** p < 0.01, *** p < 0.001.

### Increased proportion of Treg among heart-infiltrating CD4^+^ T cells after CD28-SA infusion

In infarcted mice, 4D12 treatment led to an increase in the proportion of Treg among CD4^+^ T cells of mediastinal lymph nodes as well as the heart (7). To test whether the latter was also true in pigs, we isolated CD45^+^ leukocytes from the infarcted myocardium to further analyse CD4^+^ T cells. As for peripheral blood, spleen and mediastinal lymph nodes there were no changes in the proportion of CD4^+^ T cells among total leukocytes after 4D12 infusion (Figure 4A). However, in contrast to blood and secondary lymphoid organs, we observed an increase in Treg among heart-infiltrating CD4^+^ T cells (Figure 4B). Like in spleen and mediastinal lymph nodes, the proportion of Ki-67^+^ cells among cardiac Treg (Figure 4C) and Tconv (Figure 4D) was not increased after 4D12 infusion.

**Figure 4.**
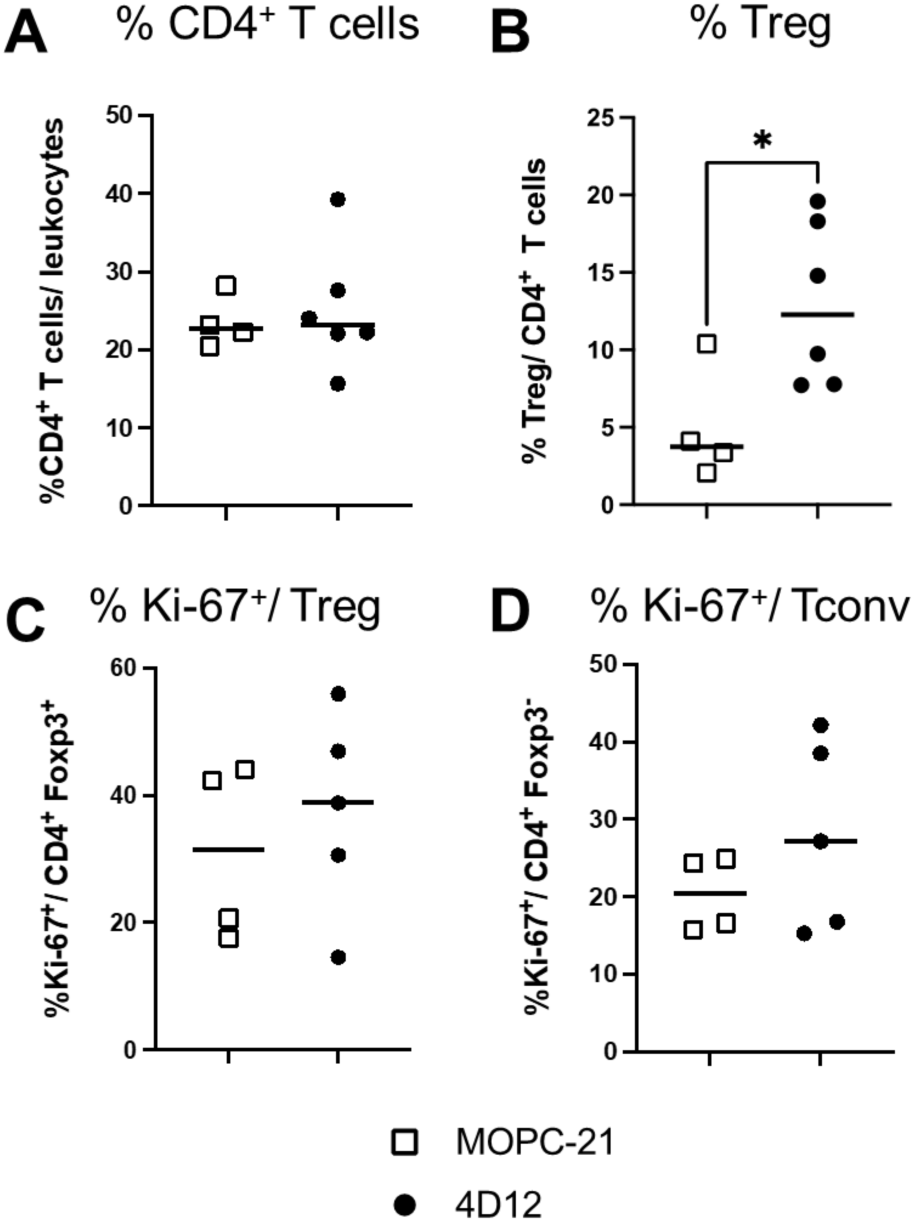
CD28-SA infusion increases the proportion of Treg among cardiac CD4^+^ T cells isolated from the infarct core. **A**) Proportion of CD4^+^ T cells among cardiac leukocytes. **B**) Frequencies of Treg among cardiac CD4^+^ T cells. **C**) Proportion of Ki-67^+^ cells among cardiac Treg and **D**) Tconv. Stats: One-tailed Mann-Whitney-Test; * p < 0.05.

To corroborate the increase in the proportion of Treg among cardiac CD4^+^ T cells after 4D12 infusion we analysed and enumerated total CD4^+^ T cells as well as CD4^+^ Foxp3^+^ double-positive Treg by immunofluorescence histology (Figure 5). Autofluorescent signals detected in the green channel allowed us to separate the infarct core from the border zone and the remote myocardium (Figure 5A). For CD4 we observed a cell surface staining, while the Foxp3 signal overlaid with the DAPI signal indicating nuclear localisation (Figure 5B). The density of CD4^+^ Tconv was much higher in the border zone and the infarct core compared to the remote myocardium irrespective of the treatment the pigs had received (Figure 5C). In MOPC-treated pigs, Treg accumulated far less, if at all, in the border zone and infarct core (Figure 5D). This is in contrast to Treg of 4D12-treated pigs, for which we observed an accumulation in the border zone and the infarct core compared to the remote myocardium (Figure 5D). Low-dose 4D12 infusion, thus, led to an increase in the proportion of Treg among cardiac CD4^+^ T cells and to an accumulation of Treg in the border zone as well as the infarct core.

**Figure 5.**
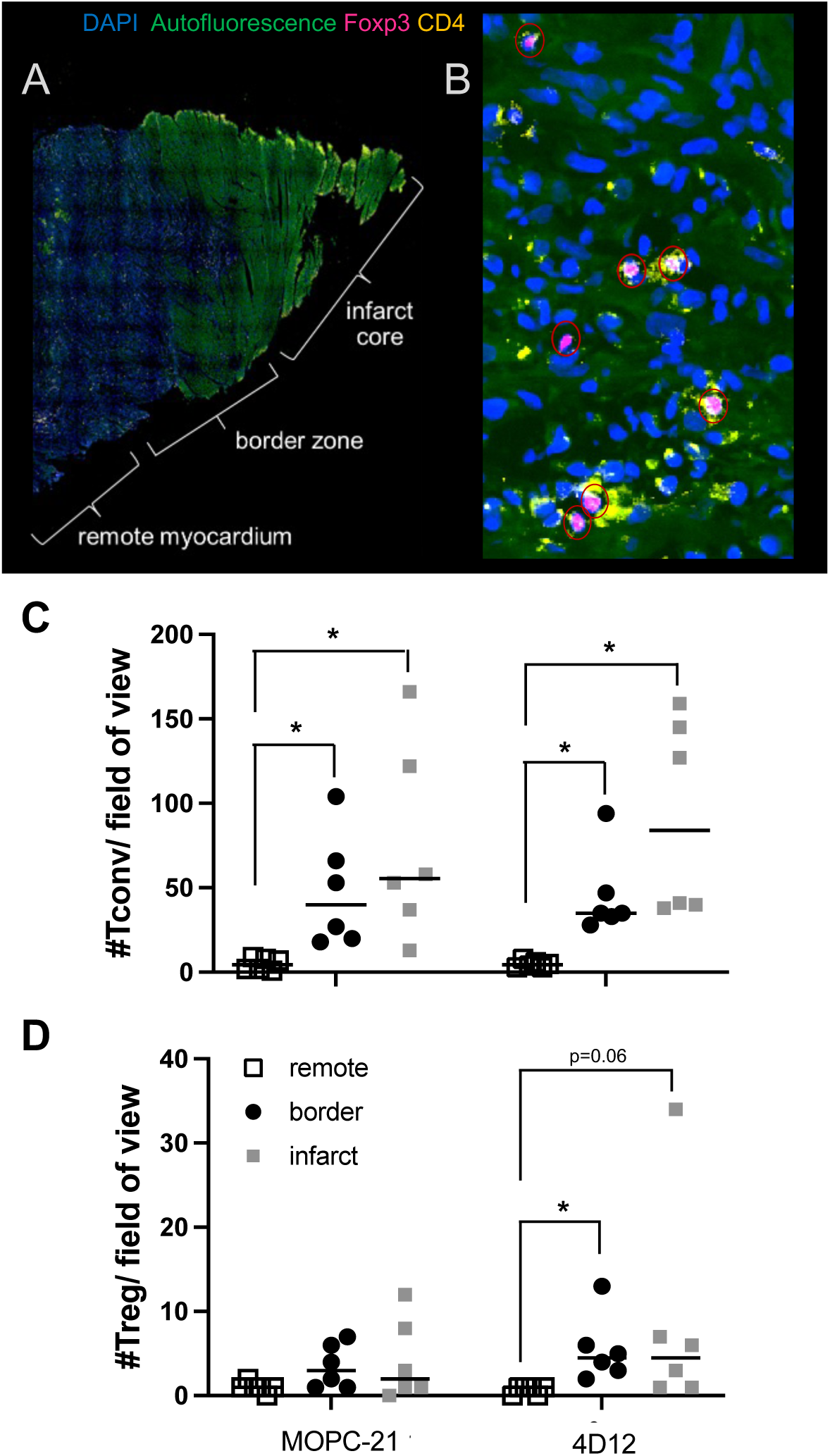
Elevated numbers of Treg in the border zone after superagonistic anti-CD28 mAb infusion. **A**) Overview showing the infarct core, the border zone and the remote myocardium seven days after MI. **B**) Detection of CD4^+^ Foxp3^−^ Tconv (yellow) and CD4^+^ Foxp3^+^ Treg (double positive – yellow and pink, highlighted by red circles) in the border zone. **C**) Tconv and **D**) Treg per field of view (715 um x 530 um). Up to six fields of view were analysed. 4D12: CD28-SA 10 µg/ kg bw; MOPC-21: control mAb. Stats: Two-tailed paired Wilcoxon test; * p < 0.05.

### CD28-SA infusion reduced the proportion of TNFα producers among cardiac myeloid cells

Treg-targeting therapies promote wound healing and prevent adverse remodelling of the myocardium by accelerating the switch of heart-infiltrating myeloid cells from a pro-inflammatory to a wound healing-promoting phenotype (17). We used TNFα production as a surrogate marker for pro-inflammatory differentiation (18) and, thus, determined the proportion of cells capable of producing TNFα among cardiac myeloid cells. 4D12 infusion did not alter the composition of the myeloid compartment in the infarcted myocardium (Figure 6A - D), but it reduced the proportion of TNFα producers among both monocytes / dendritic cells (DCs, Figure 6E) and macrophages (Figure 6F). The increase in the proportion of Treg among cardiac CD4^+^ T cells after low-dose 4D12 infusion was, thus, paralleled by a reduction in the proportion of TNFα producers among cardiac myeloid cells.

**Figure 6.**
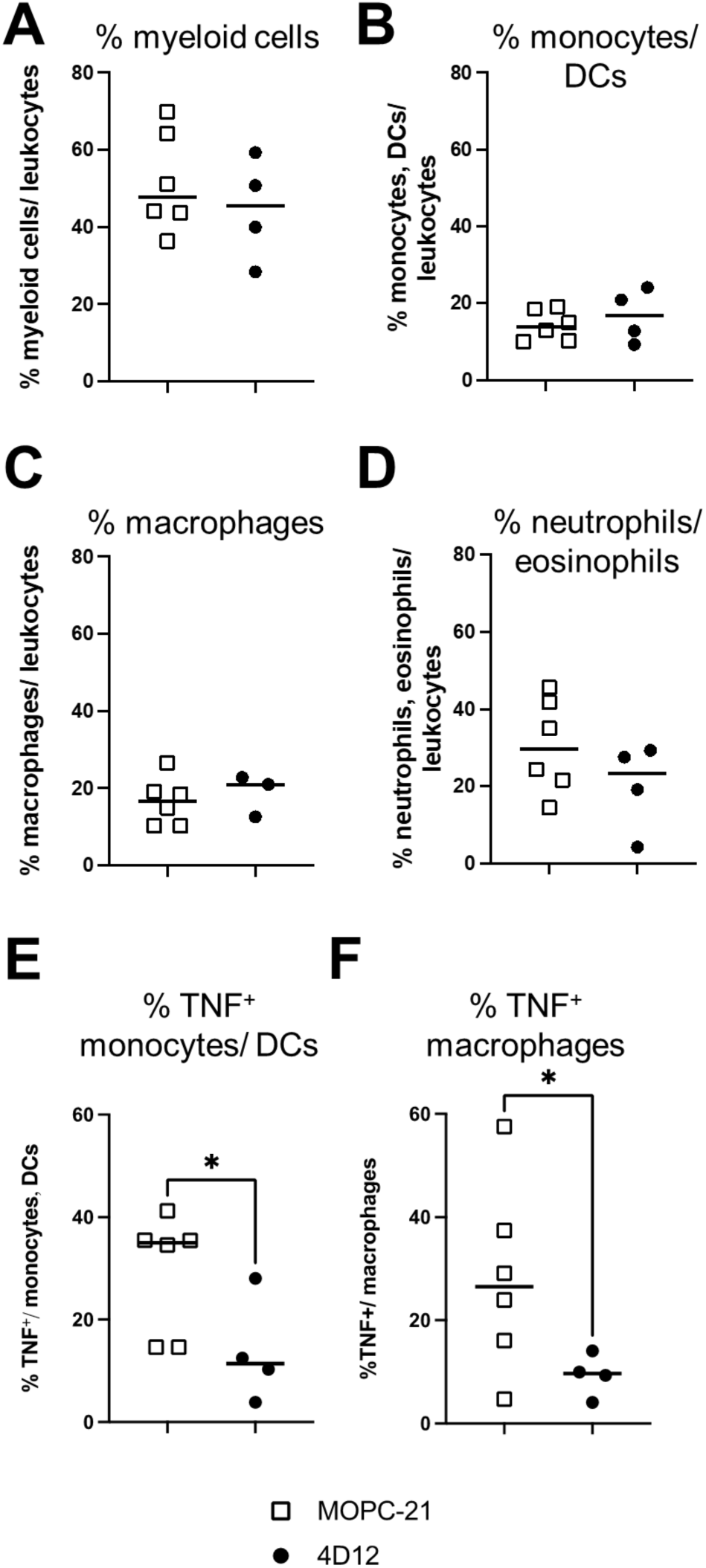
Lower proportion of TNFα- producing cells among myeloid cells following CD28-SA infusion. Proportion of **A**) cardiac myeloid cells, **B)** cardiac monocytes/DCs, **C**) cardiac macrophages, **D**) neutrophils and eosinophils among cardiac leukocytes isolated from the infarct core. Frequencies of TNFα^+^ cells after PMA/ ionomycin restimulation among **E**) monocytes/DCs and **F**) macrophages are shown. Stats: One-tailed Mann-Whitney-Test; * p < 0.05.

## Discussion

In this work we describe for the first time the short-term immunological and cardiological effects of low-dose CD28-SA infusion in pigs recovering from myocardial infarction. Our data show that low-dose CD28-SA (clone 4D12) infusion was well tolerated by the animals and did not negatively affect cardiac function during the first week after MI. The proportion of Treg was increased among heart-infiltrating CD4^+^ T cells after CD28-SA infusion with an accumulation of Treg both in the infarct core and in the border zone. Moreover, there were fewer TNFα producers among cardiac myeloid cells after low-dose CD28-SA infusion pointing towards a therapeutically enhanced switch towards better wound healing. This work is, thus, the first to show that low-dose CD28-SA treatment can be used to achieve therapeutic Treg activation in a species which is prone to develop a cytokine release syndrome after high-dose CD28-SA infusion (14).

A key observation of our study was that the proportion of Treg among CD4^+^ T cells was increased only in the infarcted myocardium. This is in contrast to mice where we observed an increase among CD4^+^ T cells of the heart and, in parallel, the mediastinal lymph nodes after MI (6, 7). Currently, it is unclear whether there are factors in the infarcted myocardium that cause this selective increase in the proportion of Treg among CD4^+^ T cells or whether Treg and Tconv receive a form of ‘licensing’ e.g. in the mediastinal lymph node after CD28-SA infusion, influencing their migratory pattern that then leads to the increase in Treg frequencies among cardiac CD4^+^ T cells. Furthermore, timing of both therapeutic applications and analyses could also contribute to the differences as the course of the immune response in the heart differs between pigs and mice (15).

Differences in migration might also account for the accumulation of Treg in the border zone and the infarct core on day seven after MI induction and four days after CD28-SA infusion. By this time point, the majority of myeloid cells have already migrated further into the infarct core (15). Thus, Treg in the border zone may modulate inflammation in the infarct core by releasing soluble mediators such as IL-10 or TGF-β. Alternatively, direct contact between these Treg and myeloid cells en route to the core may have lowered the myeloid cells’ capacity to produce TNF-α. A third possibility is that only Treg in the infarct core are biologically active, for which, however, there is currently no evidence.

Assuming that low-dose CD28-SA application will in the future be assessed in patients after MI to improve long-term outcome, the data from the pig model provide information on how low-dose CD28-SA infusion could be monitored. On the one hand, serial blood draws before and after CD28-SA infusion could be used to detect CD28-SA bound to CD28 expressed by circulating T cells (Figure 2). On the other hand, upregulation of CD28 expression on circulating T cells as well as induction of proliferation in Tconv cells (Figure 2) could be used as further biomarkers to monitor the activity of the CD28-SA *in vivo*. Mechanistically, the upregulation of CD28 on circulating as well as splenic and lymph node-CD4^+^ T cells means that enhanced CD28 co-stimulation can be expected beyond the activity of the CD28-SA itself. Further comparing signals coming from CD28-SA bound to CD28 after *in vivo* infusion to overall CD28 expression levels (Figure 2, 3), confirmed low receptor occupancy, i.e. truly low-dose infusion. An alternative to low-dose CD28-SA infusion is low-dose IL-2 treatment which has been used successfully in patients to treat e.g. graft versus host disease (19) and which has also been applied in patients with ischaemic heart disease without causing severe adverse events (20). As for the CD28-SA, IL-2 infusion also comes with the risk of inducing substantial side effects when higher dosages are applied. However, the pathophysiological cascades are quite different. High-dose CD28-SA treatment induces a CRS (21), while high-dose IL-2 infusion causes capillary leak syndrome (22), which can, however, be equally life-threatening. To enhance the safety of IL-2 treatments IL-2 muteins have been developed with an even higher affinity for the trimeric high-affinity (αβγ) versus the dimeric intermediate-affinity (βγ) IL-2 receptor than wild-type IL-2 (23). Still, a subpopulation of memory Tconv cells expresses the trimeric receptor in humans under steady state conditions so that such muteins will also target these cells in addition to targeting Treg. Currently, no adapted formats of CD28-SA are available so that induction of a CRS can only be avoided by infusing a low dose of the reagent. Apart from infusing stimulatory reagents for Treg activation the Treg - Tconv balance can also be tipped towards the Treg by blocking costimulation (10). The most direct approach here is to use a non-superagonistic anti-CD28 mAb or an antibody fragment that blocks binding of the natural ligands of CD28, i.e. CD80 and CD86. With FR104 an anti-CD28 single chain Fv is available for which phase II clinical trials have been successfully completed (24). To ensure blockade of CD28 costimulation, of course, the amounts of reagent injected are about 1,000 to 10,000-fold higher than for low-dose CD28-SA therapy, depending on the size of the molecule used for blocking. This means that the production costs for these two different approaches will also greatly vary. Another possibility for blocking CD28 costimulation is to use CTLA-4-Ig (Abatacept) which binds to CD86/ CD80-expressing antigen-presenting cells (25). E.g. patients with rheumatoid arthritis are treated with CTLA-4-Ig where its clinical value is well documented (26) - despite the fact that CTLA-4-Ig also blocks the activity of the inhibitory receptor CTLA-4. Up to now, no data are available concerning the use of CTLA-4 in patients after MI.

To understand whether CD28-SA treatment also has a positive impact on cardiac function in the mid and long term we are currently conducting experiments with serial 7 Tesla cardiac MRI (27) up to week five after myocardial infarction as the main read-out. However, for now missing long-term outcome data are one of the limitations of our study. Another limitation of our study is the low number of pigs in each treatment group, which is due to the known constraints regarding large animal experiments. If the promising shift in the cardiac myeloid compartment towards better wound healing detected on day seven post-infarction is as predictive as similar changes in mice (6, 7), then we can expect low-dose CD28-SA treatment to reduce adverse remodelling and also preserve cardiac function long-term in pigs.

In humans, low-dose CD28-SA application with dosages up to 7 µg/ kg bw has been tested in healthy volunteers and was very well tolerated (28). Following this observation, a number of successive phase II clinical trials using low-dose CD28-SA application were initiated in rheumatoid arthritis, psoriasis, systemic lupus erythematosus and patients with solid neoplasms (29). While further results and also the precise dose of CD28-SA infused have not been published so far for any of these trials, it is plausible to assume that at least in the first of these trials no severe adverse events have been observed. For a phase Ib trial in patients with rheumatoid arthritis it was stated: ‘While the details of this trial are currently being prepared for publication, it can be summarised that adverse events (AEs) remained at an acceptable level (brief episodes of fever associated with circulating IL-6)’ (30). Therefore, it may very well be feasible to set up a clinical trial in patients studying the effect of low-dose CD28-SA infusion on mid- and long-term cardiac outcome. Positive results from our ongoing study in pigs monitoring function and morphology of the heart by cardiac MRI for up to five weeks would add further momentum to such a project.

## Abbreviations

A4C: 4-chaber view
AE: adverse event
BSA: bovine serum albumin
BW: bodyweight
BZ: border zone
CD28-SA: superagonistic monoclonal anti-CD28 antibody clone 4D12
CRS: cytokine release syndrome
DC: dendritic cell
DMSO: dimethyl sulfoxide
DPBS: Dulbecco’s balanced salt solution
DTAC: dodecyltrimethylammonium-chloride
E/A: mitral valve inflow profile
ECG: electrocardiography
E/É: mitral annulus velocities
FACS: fluorescence activated cell sorting
HBSS: Hankś balanced salt solution
IC: infract core
IL: interleukin
i.m.: intramuscular
i.v.: intravenous
LAD: left anterior descending artery
LVEDV: left ventricular enddiastolic volume
LV-EF: left ventricular ejection fraction
LF-FS: left ventricular fractional shortening
LVIDd: left ventricular internal diastolic diameter
LVOT: left ventricular outflow tract
mAb: monoclonal antibody
MACS: magnetic activated cell sorting
MI: myocardial infarction
M-Mode: motion mode
MOPC-21: isotype control monoclonal antibody
PBMC: peripheral blood mononuclear cells
PBS: phosphate-buffered saline
PFA: paraformaldehyde
PLAX: parasternal long-axis view
RM: remote myocardium
SV: stroke volume
TBS: tris-buffered saline
Tconv: conventional T cells
TNFα: tumor necrosis factor alpha
Treg: regulatory T cells

## Declarations

## Acknowledgements

The authors would like to thank Katja Blouin, Helga Wagner, Nicole Maier, Sandra Werner-Wittig, Franziska Barthelmes and Jutta Meißner-Weigl for their outstanding and supportive work in the laboratory as well as Verena Burkard and Kerstin Körner for their invaluable help with animal experiments and for their dedicated and loving animal care.

## Data availability

The data supporting the conclusions of this study are provided within the paper and its supplementary information files or can be accessed from the corresponding author upon reasonable request.

