## Supplementary Information for "Low-dose infusion of a superagonistic anti-CD28 monoclonal antibody ameliorates inflammation after acute myocardial infarction in pigs"

Niklas Beyersdorf, M.D.

Institute for Virology and Immunobiology

University of Würzburg

Versbacher Straße 7

97078 Würzburg, Germany

### Supplementary Information

**Supplemental Figure S1. Coronarangiography and macroscopically visible pathological changes in pig model of myocardial infarction. A**) The LAD and its first lateral branch (blue star) were visualized by coronary angiography. The (upper) image shows the vessel before and (lower) during myocardial infarction induction with an inflated occlusion balloon. **B**) The lighter infarcted area of the pig hearts can be identified macroscopically (colored line) – one example of an explanted heart of a control mAb and of a CD28-SA treated animal. **C**) A slice from the left ventricle to illustrate the sampled areas. The three zones (infarct core, border zone and remote myocardium) can be differentiated macroscopically.

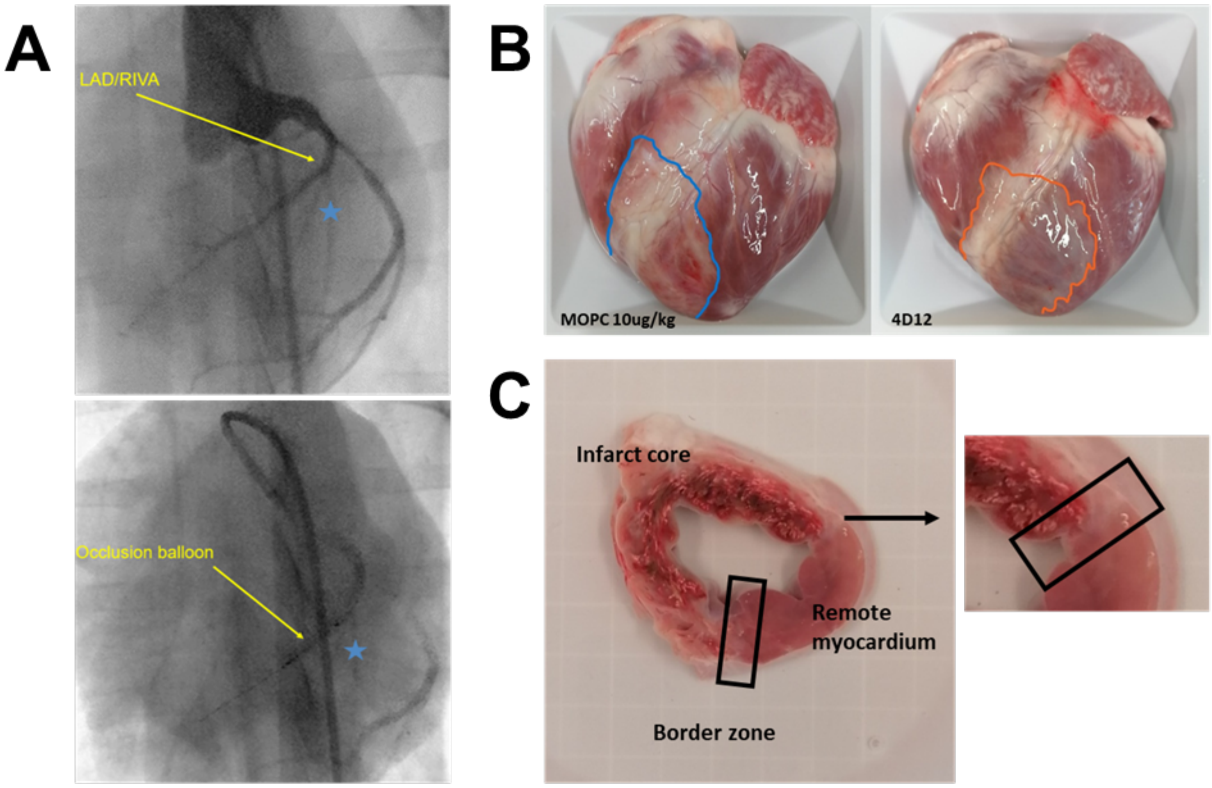

**Supplemental Figure S2.**

**Gating strategy to monitor antibody infusion and T cell subsets in peripheral blood. A**) Identification of CD4$^+$ and CD8$^+$ T cells in peripheral blood. **B**) detection of infused mAb 4D12 on CD4$^+$ T cells on day 7 (black) compared to day 0 (grey). **C**) CD28 expression on peripheral blood CD4$^+$ T cells before (grey) and seven days after MI induction (black). Dashed lines show controls without primary and secondary antibody for detection of CD28 expression. **D**) Detection of TruCount beads (left) and leukocyte gating (right) to determine absolute numbers of **E**) CD3$^+$ CD45$^+$ CD4$^+$ and CD8$^+$ T cells per ml of blood. **F**) Detection of CD25$^+$ Foxp3$^+$ Treg among gated CD4$^+$ T cells. G) Ki-67 versus Foxp3 expression of CD4$^+$ T cells of peripheral blood.

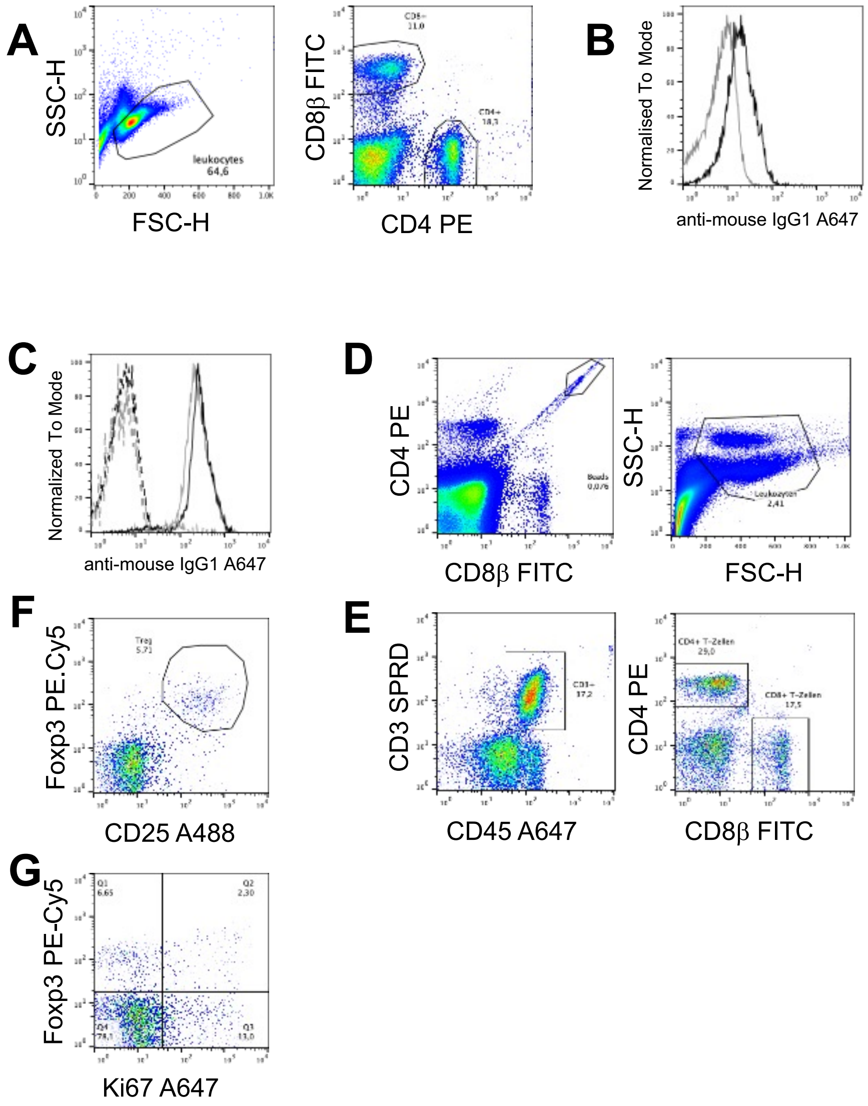

**Supplemental Figure S3.**

**Gating strategy to identify TNF-producing myeloid cells in the infarcted. A**) Identification of CD172a$^{+}$ myeloid cells. **B**) Gating of CD203a$^{+}$ macrophages. **C**) Monocytes and dendritic cells (DCs) were gated as CD203a$^{-}$ non-macrophages that were also negative for SWC8. **D**) SWC8$^{+}$ myeloid cells comprise neutrophils and eosinophils. **E**) Stimulation with PMA/ionomycin (top) increased the percentage of TNF$^{+}$ cells among the different subsets compared to cells cultured in medium (bottom).

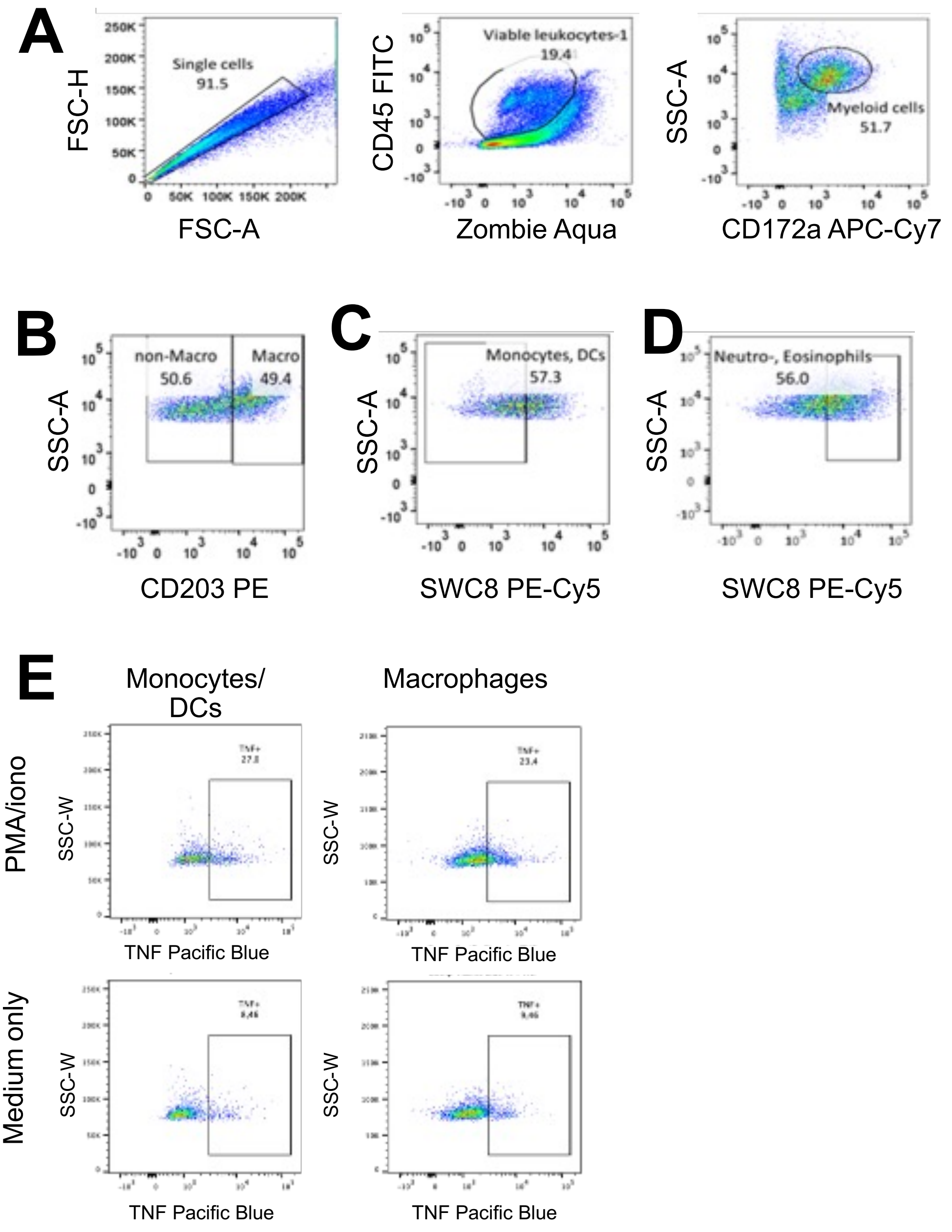

**Supplemental Video S1.**

**Echocardiographic video showing short axis view at baseline before myocardial infarction.**

https://fex.ukw.de/public/download-shares/PFhlv7A3dlmMhO7oWMB41LIXyDzL0Eyh

**Supplemental Video S2.**

**Echocardiographic video showing short axis view 7 days after myocardial infarction, showing septum akinesia.**

https://fex.ukw.de/public/download-shares/3MJypBqCG69RjYI3EDDXFV13EG0tmfYE

**Supplemental Video S3.**

**Echocardiographic video showing 4-chamber view at baseline before myocardial infarction.**

https://fex.ukw.de/public/download-shares/3MJypBqCG69RjYI3EDDXFV13EG0tmfYE

**Supplemental Video S4.**

**Echocardiographic video showing 4-chamber view 7 days after myocardial infarction showing septum akinesia.**

https://fex.ukw.de/public/download-shares/LR9h1M0Wl4Qio41zsQuaxrc9cJzurDpc

### Supplemental Table 1

**List of monoclonal primary antibodies used for flow cytometry (FC) analysis of whole blood samples**

| Antigen | Conjugate | Isotype | Clone | Species | Format | Source | Concentration |
|---|---|---|---|---|---|---|---|
| CD3ε | SPRD | IgG1 | PPT3 | mouse:pig | labeled | southern biotech | 1:500 |
| CD4α | PE | IgG2b | 74-12-4 | mouse:pig | labeled | BD | 1:100 |
| CD8β | FITC | IgG1 | PPT23 | mouse:pig | labeled | Biorad | 1:400 |
| CD45 | A647 | IgG1 | K252.1E4 | mouse:pig | labeled | Biorad | 1:15 |

### Supplemental Table 2

**List of monoclonal primary antibodies used for flow cytometry (FC) analysis of PBMC, spleen, lymphnode, thymus, heart samples**

| Antigen | Conjugate | Isotype | Clone | Reactivity | Format | Source | Concentration |
|---|---|---|---|---|---|---|---|
| unknown | / | all | polyclonal | mouse | unlabelled | Sigma Aldrich | 1:50 |
| pig CD4α | PE | IgG2b | 74-12-4 | mouse:pig | labelled | BD | 1:100 |
| pig CD8β | FITC | IgG1 | PPT23 | mouse:pig | labelled | Biorad | 1:400 |
| Foxp3 | PE-Cy5 | IgG2a | FJK-16s | rat:mouse | labelled | Thermo Fisher | 1:100 |
| Ki67 | A647 | IgG1 | B56 (RUO) | mouse:human | labelled | BD | 1:300 |
| pig CD25 | A488 | IgG1 | K231.3B2 | mouse:pig | labelled | Biorad | 1:200 |
| mouse IgG | A647 | IgG1 | RM61-1 | rat:mouse | labelled | BioLegend | 1:2000 |
| CD28 | | | 3D11 | | | in vivo Biotech | 1:195 |
| CD28 | | | 4D12 | | | in vivo Biotech | 1:105 |

### Supplemental Table 3

#### List of monoclonal primary antibodies used for flow cytometry (FC) analysis and analysis of cytokines

| Antigen | Conjugate | Isotype | Clone | Species | Format | Source | Concentration |
|---|---|---|---|---|---|---|---|
| TNF-α | Pacific Blue | IgG2 | MAb11 | mouse:human | labeled | BioLegend | 1:100 |
| SWC8´ | none | IgM | MIL3 | mouse:pig | unlabeled | Thermo Fisher | 1:20 |
| mIgM (anti SWC8) | PE-Cy5 | IgG2a | II/41 | rat:mouse | labeled | Thermo Fisher | 1:200 |
| CD203α | PE | IgG1 | PM18-7 | mouse:pig | x | Biorad | 1:200 |
| ZA | Alexa Fluor 780 | / | / | / | / | BioLegend | 1:200 |
| CD172α | APC-Cy7 | IgG2b | 74-22-15a | mouse:pig | x | BD | 1:200 |

x = Unlabeled antibody, self-conjugated using a Lightning-Link kit per manufacturer's instructions (Abcam, UK)

### Supplemental Table 4

#### Flow cytometry staining workflow

| Row / Panel | 20 min on ice | Wash 1 | 15 min on ice | Wash 2 | 15 min on ice | 15 min on ice | Wash 3 | 30 min on ice | Wash 4 | 45 min at RT (in the dark) | Wash 5 |
|---|---|---|---|---|---|---|---|---|---|---|---|
| **1** | --- | 1x, then transfer to the wells; 1x | amIgA647 | 1x | normal mouse Ig (nmIgG) | CD8β-FITC CD4-PE | 2x | | | | |
| **2** | 4D12 | 1x, then transfer to the wells; 1x | amIgA647 | 1x | normal mouse Ig (nmIgG) | CD8β-FITC CD4-PE | 2x | | | | |
| **3** | | | | 1x | normal mouse Ig (nmIgG) | CD8β-FITC CD4-PE | 2x | | | | |
| **4** | | | | 1x | normal mouse Ig (nmIgG) | CD8β-FITC CD4-PE CD27-APC | 2x | | | | |
| **5** | | | | 1x | normal mouse Ig (nmIgG) | CD25 A488 CD4 PE | 1x PBS | Fix/Perm | 1x | FoxP3 PE-Cy5 Ki67 A647 | 2x |
| **6** | | | | 1x | normal mouse Ig (nmIgG) | CD4-PE | 1x PBS | Fix/Perm | 1x | FoxP3 PE-Cy5 Ki67 A647 | 2x |
| **7** | | | | 1x | normal mouse Ig (nmIgG) | CD4-PE | 1x PBS | Fix/Perm | 1x | --- | 2x |
| | 50 µl | 150 µl FACS-buffer | 50 µl | 150 µl FACS-buffer | 25 µl | | 150 µl | 100 µl | 100 µl PermP | 50 µl | 150 µl FACS-Puffer |
